# Potential of Korean Forest Tree Seed Extracts as Multifunctional Bioresources: Evaluation of Antioxidant, Anti-inflammatory, Whitening, and Anticancer Activities

**DOI:** 10.64898/2026.03.12.711400

**Authors:** Hamin Lee, Kyungtae Park, Bo-Kook Jang, Yong-Rak Kwon, Ju-Sung Cho

## Abstract

Forest tree seeds are mass produced for afforestation and forest restoration programs, but are mostly underutilized beyond propagation. Here, we aimed to evaluate the antioxidant, anti-inflammatory, anticancer, and tyrosinase-inhibitory activities of seed extracts of seven economically important forest tree species in the Republic of Korea to explore their potential as multifunctional natural bioresources. The seed extracts of *Alnus japonica, Chamaecyparis obtusa, Cornus kousa, Phellodendron amurense, Pinus densiflora, Prunus sargentii*, and *Quercus glauca* were comparatively assessed using multiple *in vitro* assays. The results revealed clear species-dependent functional profiles rather than uniform bioactivities across species. *Quercus glauca* exhibited strong antioxidant activity and significant anti-inflammatory and tyrosinase-inhibitory activities, suggesting multifunctional potential, while *C. obtuse* presented considerable anticancer activity against several cancer cell lines. *Alnus japonica* exhibited the highest tyrosinase-inhibitory activity, followed by *Q. glauca* and *C. obtuse*; *A. japonica* extract also showed a strong antioxidant capacity. Overall, the results demonstrated that forest tree seed extracts possess diverse and complementary bioactivities, supporting their potential as underexplored multifunctional natural materials. By focusing on seed resources generated within existing afforestation systems, we highlight a sustainable approach to valorize forest-derived by-products without additional pressure on natural ecosystems. Nevertheless, as bioactivities were evaluated using crude extracts, further studies are required to identify and elucidate the active compounds and their mechanisms of action.

## Introduction

South Korea is highly forested, with approximately 63% of its national land area covered by forests, supporting 715 woody plant species among 3,960 native plant species. The forest ecosystems function as not only major reservoirs of biodiversity but also important sources of plant genetic resources with potential applications across pharmaceutical, food, cosmetic, and bio-based industries [1,2]. Forest plants shaped by distinct climatic and edaphic conditions often exhibit unique secondary metabolite profiles and have long been utilized in traditional medicine, functional foods, and natural health products [3]. In recent years, the rapid expansion of plant-based industries, driven by increasing consumer preference for sustainable and “clean-label” products over synthetic alternatives, has markedly intensified demand for natural-origin raw materials [4–7]. Simultaneously, growing industrial demand for plant resources has raised concerns regarding overharvesting, ecosystem degradation, and accelerated biodiversity loss [8]. In this milieu, the seed orchard system of the Republic of Korea represents a structural system capable of mitigating such ecological risks [9]. Unlike spontaneous or small-scale cultivation approaches that may rely on initial wild harvesting, South Korean seed orchards are well-established, with state-managed facilities consistently producing genetically standardized, high-quality seeds in bulk. This organized and traceable production system discourages illegal wild collection and enables industrial utilization without exerting additional pressure on natural forest ecosystems, thereby meeting increasing demand while safeguarding forest resources.

Forest tree seeds represent a particularly underexplored yet promising bioresource. While seeds are primarily produced and used for afforestation and propagation, surplus seeds or those excluded during quality refinement and long-term storage are frequently discarded or remain underutilized from an industrial perspective. Here, we focused on the following species: *Pinus densiflora, Chamaecyparis obtusa*, and *Quercus glauca* classified as timber species; *Alnus japonica* and *Prunus sargentii* as pollution-tolerant species; *Cornus kousa* as a landscaping species; and *Phellodendron amurense* as a fire-resistant species. All of these species are designated as “recommended afforestation species” by the Korea Forest Service [10], ensuring stable and large-scale seed production without imposing additional ecological burdens.

Beyond their silvicultural roles, the aforementioned species have been reported to exhibit diverse biological activities. For instance, the bark and leaves of *P. amurense* possess anti-inflammatory and antiproliferative properties [11,12], while essential oils derived from *P. densiflora* and *C. obtusa* demonstrate antimicrobial and antiviral activities [13–15]. *Quercus* species contain a wide array of phytochemicals, including phenolic acids, tannins, flavonoids, and triterpenes, conferring high industrial potential [16]. Additionally, antioxidant and antimicrobial activities of the leaves and bark of *P. sargentii* [17], anti-elastase activity of the stems of *C. kousa* [18], and antifungal and antiparasitic effects of the bark and stems of *A. japonica* [19] has been reported.

Forest tree seeds have received relatively limited attention in terms of their biochemical composition and functional bioactivities. Seeds are metabolically distinct organs with high concentrations of phenolic compounds, antioxidants, and defense-related metabolites. However, the potential of seeds, particularly those of Korean forest tree species, as functional bioresources remains poorly characterized. The utilization of surplus or non-viable seeds generated as by-products of existing afforestation systems offers a practical opportunity to enhance resource-use efficiency without intensifying harvesting pressure on wild populations. Therefore, in the present study, we aimed to comprehensively evaluate the antioxidant, anti-inflammatory, whitening, and anticancer activities of seed extracts of seven forest tree species widely used in South Korean afforestation programs. By elucidating the functional potential of these underutilized seed resources, we intend to contribute to the development of sustainable plant-based industries.

## Materials and Methods

### Plant materials and seed collection

The seeds of seven forest tree species—*A. japonica, C. obtusa, C. kousa, P. amurense, P. densiflora, P. sargentii*, and *Q. glauca*—were obtained from the National Forest Seed and Variety Center (NFSV; Chungju, Korea). All seeds were harvested between 2017 and 2023 from state-managed seed orchards located in various regions of South Korea. Until extraction, the seeds were stored under low-temperature, dry conditions (4°C). Table 1 provides the scientific names of these species, their family names, production sites, and quality metrics. Here, only seed lots classified as preliminary grade under the NFSV quality control standards were used. According to this classification, seeds are excluded from national afforestation and propagation programs if their germination rate or efficiency values fall below the standardized thresholds for forest restoration.

**Table 1.** List of forest trees species used.

| Scientific name | Family name | Abbreviation<sup>z</sup> | Year of production | Production site | Germination rate (%) | Efficiency (%) | Seed quality |
| --- | --- | --- | --- | --- | --- | --- | --- |
| *Alnus japonica* | Betulaceae | AJ | 2018 | Jeongseon-gun, Gangwon-do | 10.5 | 8.0 | Preliminary |
| *Chamaecyparis obtuse* | Cupressaceae | CO | 2023 | Chungju-si, Chungcheongbuk-do | 10 | 20 | Preliminary |
| *Cornus kousa* | Cornaceae | CK | 2020 | Muju-gun, Jeollabuk-do | 5.5 | 5.5 | Preliminary |
| *Phellodendron amurense* | Rutaceae | PA | 2021 | Hongcheon-gun, Gangwon-do | 0 | 0 | Preliminary |
| *Pinus densiflora* | Pinaceae | PD | 2017 | Hongcheon-gun, Gangwon-do | 80.1 | 25.2 | Preliminary |
| *Prunus sargentii* | Rosaceae | PS | 2021 | Yanggu-gun, Gangwon-do | 0 | 0 | Preliminary |
| *Quercus glauca* | Fagaceae | QG | 2021 | Seogwipo-si, Jeju-do | 36.5 | 2.0 | Preliminary |

<sup>z</sup>Species abbreviations are used in the figures and text for clarity

### Extraction process

The seeds of the seven species were freeze-dried and ground using a milling machine. For extraction, 1 g of seed powder was mixed with 50 mL of 70% ethanol, followed by ultrasonic extraction for 30 min. The extracts were used to analyze the total polyphenol and flavonoid content (TPC and TFC, respectively) and antioxidant activity. For the evaluation of anti-inflammatory, anticancer, and skin-whitening activities, the extracts were prepared as follows: after solvent removal using a rotary vacuum evaporator (N-1000-SW, EYELA, Tokyo, Japan), the residues were re-dissolved in dimethyl sulfoxide (DMSO) at a concentration of 100 mg·mL^−1^ and passed through a 0.45-μm syringe filter.

### Determination of TPC and TFC

TFC was determined using the diethylene glycol (DEG)-sodium hydroxide (NaOH) colorimetric method described by Davis [20]. A reaction mixture consisting of 200 μL of the extract, 2 mL of DEG, and 200 μL of 1 N NaOH was prepared and incubated in a water bath at 37°C for 1 h. Thereafter, the absorbance was measured at 420 nm using a spectrophotometer. A standard calibration curve was constructed using quercetin, and the TFC is expressed as quercetin equivalents (QE) per gram of dry weight (QE·mg g^−1^ DW).

TPC was determined using a modified Folin–Denis method [21]. A 100-μL aliquot of the extract was mixed with 2 mL of 2% sodium carbonate solution and allowed to react for 3 min, and then 100 μL of 50% Folin–Ciocalteu reagent was added. The mixture was incubated in the dark for 30 min, and the absorbance was measured at 750 nm using a spectrophotometer. A standard curve was prepared using gallic acid, and TPC is expressed as milligrams of gallic acid equivalents (GAE) per gram of dry weight.

### Determination of antioxidant activities

2,2′-Azino-bis(3-ethylbenzothiazoline-6-sulfonic acid) (ABTS) radical scavenging activity was determined using a modified version of the decolorization method described by Re et al. [22]. To generate ABTS^+^radicals, 50 mL of ABTS solution was mixed with 12.31 mL of 2.6 mM potassium persulfate and stirred in the dark. The resulting solution was then diluted with phosphate-buffered saline to an absorbance of 0.70 ± 0.03 at 732 nm. For the assay, 950 μL of the diluted ABTS solution was mixed with 50 μL of the extract and incubated in the dark for 10 min. The ABTS radical scavenging activity is expressed as electron donating ability (EDA, %), and the concentration of the extract required to reduce 50% of ABTS radicals (RC_50_, mg·mL^−1^) was calculated as follows:

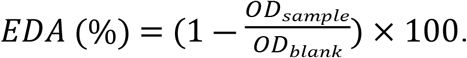

2,2-Diphenyl-1-picrylhydrazyl (DPPH) radical scavenging activity was evaluated following the method described by Brand-Williams et al. [23]. A 200-μL aliquot of the extract was mixed with 800 μL of 0.2 mM DPPH solution, and the solution was incubated in the dark for 30 min. The absorbance was then measured at 517 nm. The radical scavenging activity was calculated using the same formula that was used in the ABTS assay.

### Determination of anti-inflammatory activity

RAW264.7 cells were obtained from the Korea Cell Line Bank (Seoul, South Korea). For cell culture, Dulbecco’s modified Eagle’s medium (DMEM) supplemented with 10% fetal bovine serum (FBS), 100 units·mL^−1^ penicillin, and 100 μg·mL^−1^ streptomycin was used. The cells were sub-cultured every 2 days in an incubator maintained at 37°C with 5% CO_2_ and 95% humidity.

Cell viability was evaluated using the 3-(4,5-dimethylthiazol-2-yl)-2,5-diphenyltetrazolium bromide (MTT) assay. RAW264.7 cells were seeded in 96-well plates at a density of 2.5 × 10^5^ cells·well^−1^ and incubated for 6 h. Thereafter, 200 μL of each treatment agent was added to the wells: lipopolysaccharide (LPS, 1 μg·mL^−1^), quercetin (positive control, 25 μM), and each extract (100 μg·mL^−1^). Following 18 h of treatment, 10 μL of MTT solution (1 mg·mL^−1^) was added to each well, and the samples were incubated for an additional 90 min. The medium was then removed, and the resulting formazan crystals were dissolved in 200 μL of DMSO. Absorbance was measured at 550 nm using a microplate reader (Epoch, BioTek, Vermont, USA). Cell viability is expressed as percentage relative to that of the untreated control. Based on the criteria described in ISO 10993-5 (2009), treatments that resulted in cell viability below 80% were considered cytotoxic and excluded from further analysis.

Nitric oxide (NO) production was evaluated using the Griess assay [24]. Following cell seeding and sample treatment as described in the MTT assay, 100 μL of the cell culture supernatant was mixed with an equal volume of Griess reagent and incubated for 10 min. Thereafter, absorbance was measured at 550 nm. NO concentration was calculated using a standard curve constructed with sodium nitrate.

### Determination of anticancer activity

Six human-derived cancer cell lines were used: A549 (lung cancer), Caco-2 and HCT15 (colorectal cancer), LNCap-LN3 (prostate cancer), MDA-MB-231 (breast cancer), and SK-OV-3 (ovarian cancer). All cell lines were obtained from the Korea Cell Line Bank. The cells were cultured in RPMI 1640 medium supplemented with 10% FBS and 1% penicillin–streptomycin. All cell lines were subcultured every 2 days under standard incubation conditions of 37°C, 5% CO_2_, and 95% humidity.

To evaluate the anticancer activity of the seven forest tree seed extracts, MTT assays were performed. The six cancer cell lines were seeded in 96-well plates at a density of 1 × 10^4^ cells·well^−1^ and incubated for 24 h. The cells were exposed to one of the following three treatments and incubated for an additional 48 h: control (no treatment), mock control (treated with DMSO at the same concentration as that in the sample), and extract treatment (100 µg·mL^−1^). Thereafter, 1 mg·mL^−1^ MTT solution was added to each well, and the samples were incubated for 4 h. The medium was then removed, and the resulting formazan crystals were completely dissolved in DMSO. Absorbance was measured at 550 nm, and the relative cell viability of the extract-treated group was calculated relative to that of the mock control group.

### Determination of the tyrosinase-inhibitory activity for skin whitening

The whitening effect of the extracts was evaluated using an L-DOPA oxidation-based tyrosinase inhibition assay. Each extract (100 mg·mL^−1^ in DMSO) was diluted with 0.175 M sodium phosphate buffer (pH 6.8) to a final concentration of 1 mg·mL^−1^. To assess the effect of DMSO contained in the extracts, an equivalent volume of DMSO diluted in buffer was used as the control. Ascorbic acid (AsA, 1 mg·mL^−1^) was used as a positive control. For the assay, 20 μL of sample, 40 μL of 10 mM L-DOPA, 100 μL of buffer, and 40 μL of tyrosinase (110 units·mL^−1^) were added sequentially. The enzymatic reaction was monitored kinetically at 475 nm using a microplate reader, with absorbance measured every minute for 20 min. Tyrosinase activity (%) was calculated using the inhibition formula described by Tsai et al. [25]:

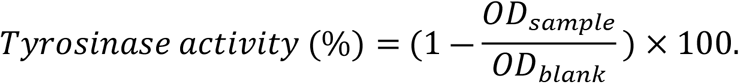

### Data collection and statistical analysis

All experimental measurements were conducted with appropriate replicates: extract yield, antioxidant activity, and tyrosinase inhibition assays were performed in triplicate, whereas anti-inflammatory and anticancer assays were conducted with six replicates. Data are presented as mean ± standard deviation (SD). Statistical analyses were conducted using the SAS software (version 9.4; SAS Institute, Cary, NC, USA). Treatment effects were evaluated using Tukey’s honest significant difference (HSD) test, with statistical significance set at *p* < 0.05.

## Results and Discussion

### Extract yield and antioxidant activities of forest tree seeds

*Quercus glauca* had the highest polyphenol content (30.7 ± 0.28 mg GAE·g^−1^), followed by *A. japonica* (25.2 ± 0.1 mg GAE·g^−1^) and *C. obtusa* (21.5 ± 0.1 mg GAE·g^−1^; Fig. 1A). Similar trends were observed for TFC (Fig. 1B), with *Q. glauca* also showing the highest flavonoid content (46.0 ± 0.25 mg QE·g^−1^). The ABTS and DPPH assays, performed to determine the radical-scavenging capacity, demonstrated that *Q. glauca* had the highest TPC and TFC and the lowest RC_50_ values, indicating the strongest antioxidant activity among all tested species (Fig. 1C, D). These results are consistent with those of Kumar et al. [26], who reported strong positive correlations between TPC and ABTS (r = 0.998, R^2^ = 0.997) and DPPH (r = 0.994, R^2^ = 0.988) radical scavenging activities of various plant extracts. The high correlation coefficients suggest that the polyphenol content is a critical contributor to antioxidant potential. The correlations observed between compound content (TPC and TFC) and radical-scavenging activities support the notion that *Q. glauca* contains abundant phenolic compounds and highly efficient radical-scavenging compounds. These results emphasize the importance of both the quantity and quality of the antioxidant constituents. The combination of high antioxidant content and radical-scavenging activity of *Q. glauca* highlights its potential as a valuable source of natural antioxidants. Furthermore, the strong correlation observed between TPC/TFC and radical-scavenging activities in *Q. glauca* suggests its potential as a premium functional forest bioresource.

**Fig 1.**
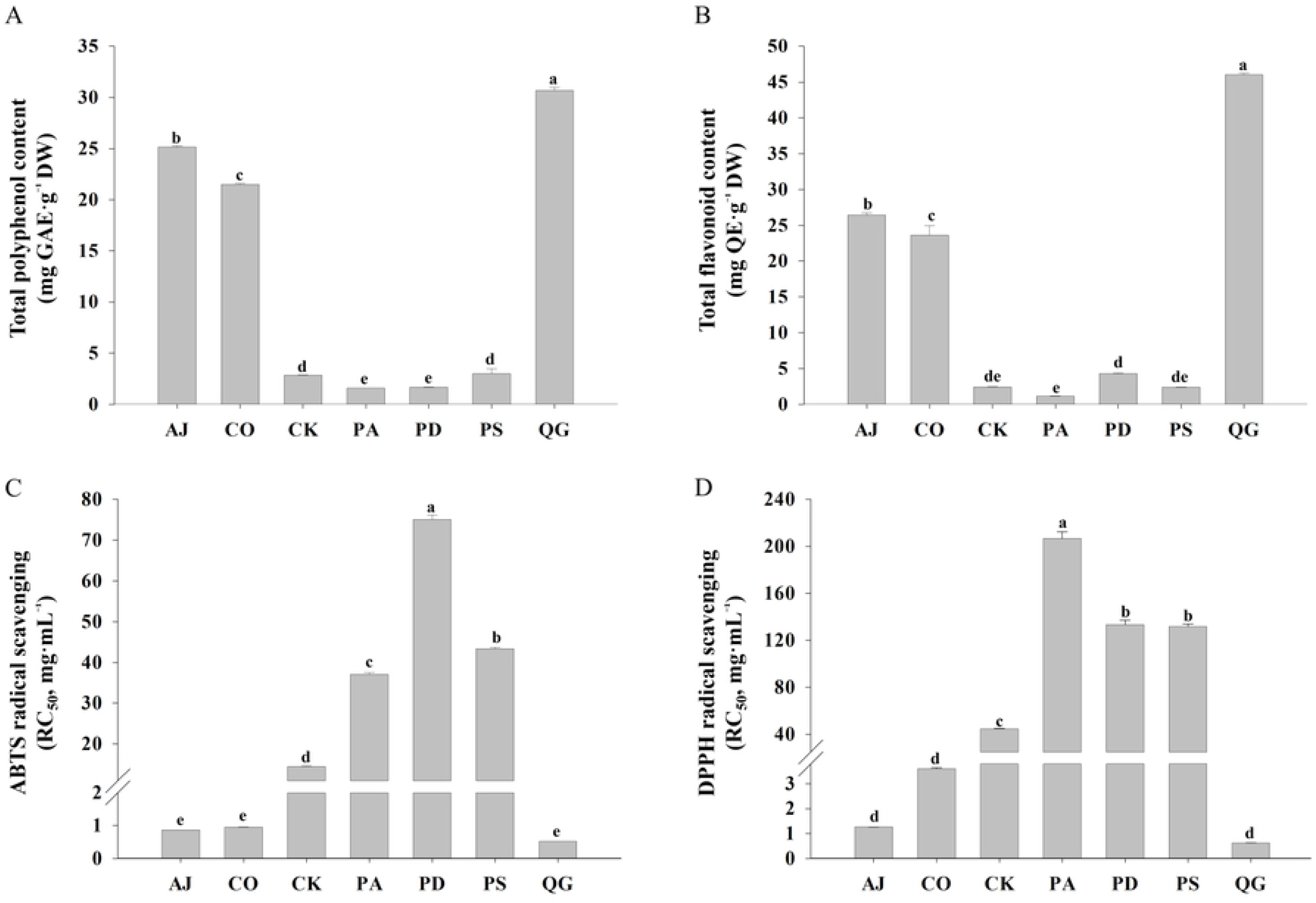
Antioxidant compound content and activities of forest seed extracts. (A) Total polyphenol content (TPC) in mg GAE·g^−1^ dry weight. (B) Total flavonoid content (TFC) in mg QE·g^−1^ dry weight. (C) ABTS radical scavenging activity. (D) DPPH radical scavenging activity. Species abbreviations (e.g., AJ, CO, and CK) are listed in Table 1. Data are presented as mean ± standard error (SE) of three replicates (n = 3). Different lowercase letters indicate significant differences among species based on Tukey’s HSD test (*p* < 0.05).

### Anti-inflammatory activity of the seed extracts on RAW 264.7 macrophages

To examine the anti-inflammatory properties of the seed extracts, we evaluated their cytotoxicity against RAW 264.7 macrophage cells. As shown in Fig. 2a, most extracts maintained cell viability above 80% at 100 µg·mL^−1^, indicating low cytotoxicity at the tested dose. However, the viability of cells treated with the seed extracts of *A. japonica* and *C. obtusa* was 77.5% and 7.0%, respectively; therefore, these extracts were classified as cytotoxic based on the ISO 10993-5 standard. Accordingly, these extracts were excluded from subsequent NO inhibition assays.

**Fig 2.**
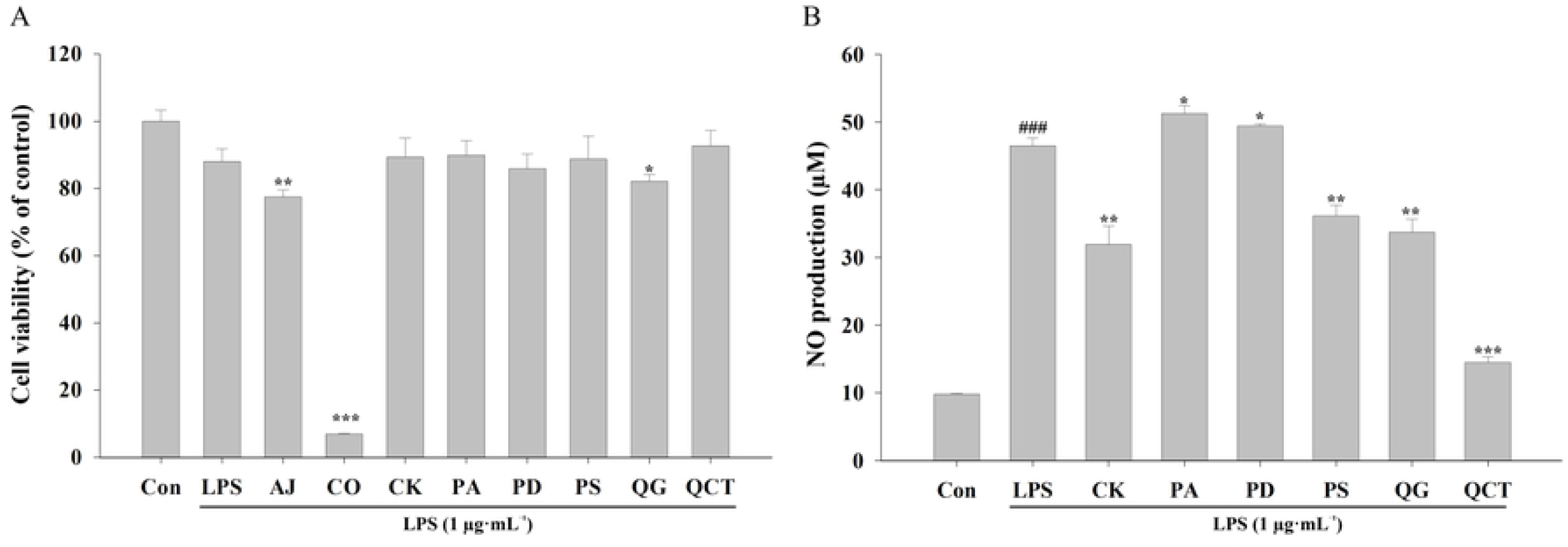
Anti-inflammatory activity of forest seed extracts in RAW 264.7 macrophages. (A) Cell viability is expressed as percentage relative to that of the control group, and significant differences among seed extract treatments were determined using Tukey’s HSD test (*p* < 0.05). (B) Nitric oxide (NO) production was measured in LPS-stimulated cells. ### and * indicate significant differences between the control and LPS (*p* < 0.001, Student’s *t*-test) and between LPS and extract-treated (\**p* < 0.05, \*\**p* < 0.01, \*\*\**p* < 0.001, Student’s *t*-test) groups, respectively. Data are presented as mean ± standard error (SE) of six replicates (n = 6). Species abbreviations (e.g., AJ, CO, and CK) are defined in Table 1.

Following LPS stimulation, NO production increased markedly to 46.51 ± 1.10 µM, compared with that in the control group (9.81 ± 0.10 µM), confirming successful induction of the inflammatory response (Fig. 2b). The positive control quercetin significantly reduced the NO level to 14.50 ± 0.84 µM. Among the seed extracts tested, *Q. glauca, C. kousa*, and *P. sargentii* extracts considerably suppressed NO production, lowering levels to 33.76 ± 1.88, 31.96 ± 2.68, and 36.16 ± 1.58 µM, respectively. These reductions were statistically significant compared with those in the LPS-treated group (*p* < 0.01). In contrast, treatment with the extracts of *P. amurense* (51.27 ± 1.18 µM) and *P. densiflora* (49.42 ± 0.33 µM) did not result in a meaningful decrease in NO production.

The above results suggest that the seed extracts of *Q. glauca, C. kousa*, and *P. sargentii* exhibit anti-inflammatory potential by effectively suppressing NO production. Numerous studies have demonstrated a strong and complex interrelationship between antioxidant activity and inflammation [27–29]. Consistent with the findings of these studies, the seed extracts of *Q. glauca, C. kousa*, and *P. sargentii* displayed pronounced anti-inflammatory activity and also exhibited significantly higher TPC, TFC, and ABTS and DPPH radical scavenging activities, indicating a significant association between antioxidant capacity and anti-inflammatory activity of these seed extracts.

Additionally, the strong anti-inflammatory activities of *Q. glauca, C. kousa*, and *P. sargentii* are consistent with those previously reported for each species. Kim et al. [30] reported that *Q. glauca* acorn shell extracts exert anti-inflammatory activity by alleviating oxidative stress and promoting recovery from hydrogen peroxide-induced cellular damage. Similarly, leaf extracts of *C. kousa* with high polyphenol content have potential benefits in the prevention and treatment of inflammatory diseases [31]. Furthermore, hexane and chloroform fractions of *P. sargentii* leaves reportedly inhibited NO production in LPS-stimulated RAW264.7 macrophages [32]. While previous studies primarily focused on leaves or bark, here we demonstrated comparable anti-inflammatory activities of seed extracts of these species. This finding suggests that multiple parts, including the seeds, of these species have anti-inflammatory potential.

### Anticancer activity against various cancer cell lines

The seed extracts exhibited varying degrees of anticancer activity against seven human cancer cell lines: lung (A549), skin (B16F10), colorectal (Caco-2 and HCT15), prostate (LNCaP-LN3), breast (MDA-MB-231), and ovarian (SK-OV-3) (Fig. 3). Among these extracts, *C. obtusa* extract demonstrated the most potent cytotoxic effect, significantly reducing the viability of multiple cancer cell lines. It reduced the viability of A549, B16F10, Caco-2, and HCT15 cells to less than 10%, suggesting strong broad-spectrum antiproliferative effects. LNCaP-LN3 and MDA-MB-231 cells also exhibited a significant reduction in viability, further supporting the potent anticancer potential of *C. obtusa* extract. In contrast, the effect on SK-OV-3 cells was relatively moderate, suggesting cell-line– specific sensitivity of the extract. In addition to *C. obtuse* seed extract, other seed extracts exhibited distinct cell-line–specific anticancer activities. *Alnus japonica* significantly inhibited the proliferation of A549, LNCaP-LN3, and HCT15 cells, reducing cell viability to 76.6% (*p* < 0.01), 35.1% (*p* < 0.001), and 78.0% (*p* < 0.001), respectively, compared to the vehicle control. However, no significant effects were observed in B16F10, Caco-2, or MDA-MB-231 cells, while a stimulatory effect on cell proliferation was detected in SK-OV-3 cells (*p* < 0.01). *Cornus kousa* significantly suppressed cell proliferation in six cancer cell lines, except in SK-OV-3. *Prunus amurense* exhibited anticancer activity against B16F10 (*p* < 0.05) and LNCaP-LN3 cells (*p* < 0.001). *Pinus densiflora* exhibited significant inhibitory effects on A549 (*p* < 0.05), LNCaP-LN3 (*p* < 0.01), and HCT15 cells (*p* < 0.001). *Prunus sargentii* demonstrated antiproliferative activity in Caco-2 and HCT15 cells (both *p* < 0.05), while *Q. glauca* significantly reduced the viability of A549 (*p* < 0.01), Caco-2 (*p* < 0.05), and LNCaP-LN3 cells (*p* < 0.001).

**Fig 3.**
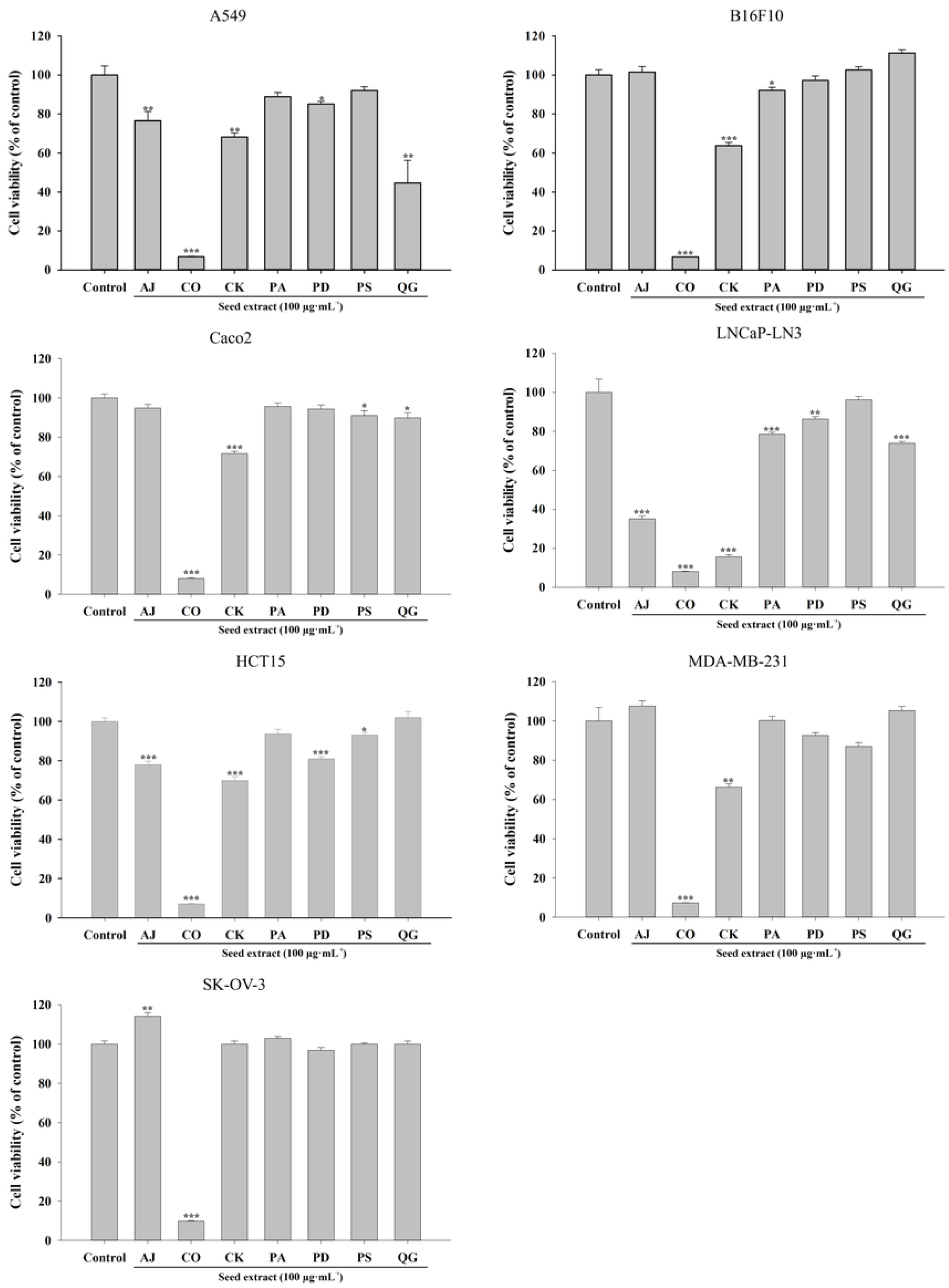
Anticancer activity of forest seed extracts on various cancer cell lines. Cell viability was measured using the MTT assay in seven human cancer cell lines: lung cancer (A549), melanoma (B16F10), ovarian cancer (SK-OV-3), colon cancer (Caco-2, HCT15), prostate cancer (LNCaP-LN3), and breast cancer (MDA-MB-231). Data are presented as mean ± standard error (SE) of three replicates (n = 6). Significant differences between the control and extract-treated groups were determined using Student’s *t*-test (\**p* < 0.05, \*\**p* < 0.01, \*\*\**p* < 0.001). Species abbreviations (e.g., AJ, CO, and CK) are defined in Table 1.

Our results demonstrate that the anticancer activities of forest tree seed extracts range from broad-spectrum effects to cell-line–specific responses. The seed extract of *C. obtusa* exhibited the most pronounced and broad-spectrum antiproliferative activity, markedly reducing cell viability across cancer cell lines of diverse tissue origins. Similarly, previous studies have reported that extracts of the leaves and heartwood of *C. obtusa* significantly inhibit the proliferation of various cancer cell lines, including PANC-1, HCT116, KB, HONE-1, and TSCH cells [33–35]. Such potent anticancer activity has been attributed to the combined effects of diverse phytochemicals abundant in coniferous species, including terpenoids, flavonoids, and phenolic acids [36]. Accordingly, the effects observed here may also be driven by similar phytochemical compositions in the seed extracts. Additionally, six seed extracts exhibited selective inhibitory effects across cancer cell lines, confirming that anticancer activity varies with both plant species and cancer cell line type. Cell-line–specific sensitivity of plant extracts has been reported [37], and such selective responses are presumed to arise from variations in the bioactive compounds in each plant species and their distinct mechanisms of action across different cancer cell lines [38]. Generally, species– and cell-line–specific responses are observed in anticancer research on plant extract; therefore, these findings highlight the importance of systematic screening involving diverse plant species and multiple cancer cell lines to accurately evaluate anticancer potential.

### Tyrosinase-inhibitory activity

The tyrosinase-inhibitory activity of the seed extracts was evaluated to assess their potential as natural skin-whitening agents. As shown in Fig. 4 and the accompanying quantitative data, all extracts exhibited varying degrees of tyrosinase activity inhibition, ranging from 28.48% to 58.90%. Ascorbic acid, used as the positive control, demonstrated the highest inhibition at 98.31% ± 0.15%, validating the assay.

**Fig 4.**
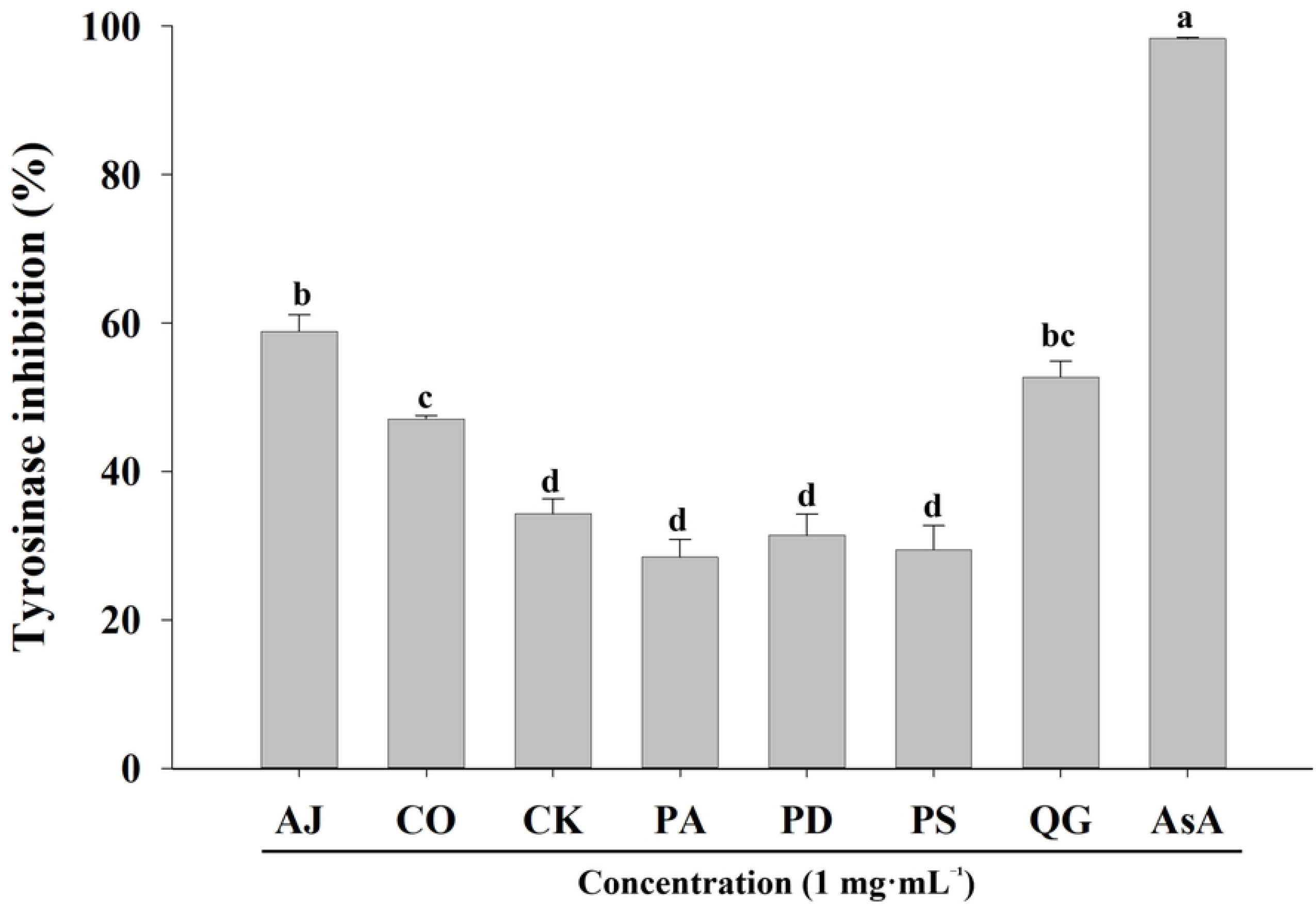
Tyrosinase-inhibitory activity of forest tree seed extracts. The inhibitory effect of seed extracts on tyrosinase activity is expressed as percentage. Different lowercase letters indicate significant differences among seed extracts based on Tukey’s HSD test (*p* < 0.05, n = 3). AsA (ascorbic acid, 1 mg·mL^−1^) was used as a positive control. Species abbreviations (e.g., AJ, CO, and CK) are defined in Table 1.

Among the extracts, *A. japonica* seed extract exhibited the strongest tyrosinase-inhibitory effect (58.90% ± 2.27%), followed by *Q. glauca* (52.75% ± 2.12%) and *C. obtusa* seed extracts (47.09% ± 0.49%). These three species exhibited significantly higher activity than the other species (*p* < 0.05). In contrast, *P. amurense, P. densiflora*, and *P. sargentii* exhibited relatively low inhibition rates (28.48–31.39%), indicating limited effectiveness in tyrosinase activity inhibition.

The aforementioned species-dependent patterns suggest that qualitative and quantitative differences in phenolic compounds and other secondary metabolites may underlie the observed variations in tyrosinase-inhibitory activity. Various phenolic compounds, including flavonoids, phenolic acids, and tannins, are known to inhibit tyrosinase activity through multiple mechanisms: (i) direct interaction with the catalytic pocket of the enzyme, (ii) chelation of the two copper ions at the active site, and (iii) acting as alternative substrates or redox competitors altering L-DOPA oxidation [39–41]. Here, the seed extracts of *A. japonica, C. obtusa*, and *Q. glauca* exhibited strong antioxidant activities along with significant tyrosinase-inhibitory effects. Previous studies have reported positive correlations among TPC, TFC, tyrosinase-inhibitory activity, and antioxidant capacity [42,43].

Accordingly, the relatively high tyrosinase-inhibitory activities exhibited by the three species are likely closely associated with their elevated TPC and TFC as well as their antioxidant activities, as assessed using ABTS and DPPH assays, consistent with the findings of a previous study on plant-derived phenolic-rich extracts [44]. Nevertheless, certain limitations of the present study should be acknowledged. Individual phenolic compounds were neither identified nor quantified here, and the specific molecules responsible for tyrosinase activity inhibition, as well as their direct mechanisms of interaction with the enzyme, were not elucidated. Future studies employing chromatographic and spectrometric approaches, such as HPLC and LC–MS, combined with bioactivity-guided fractionation, will be necessary to identify the key active constituents and to clarify their roles in tyrosinase activity inhibition.

## Conclusions

The findings of this study demonstrated the potential of forest tree seeds, including preliminary-grade by-products from state-managed seed orchards, as underutilized bioresources with diverse biological activities. The major conclusions of this study are as follows. (1) Forest tree seed extracts exhibited species-dependent antioxidant, nitric oxide inhibitory, anticancer, and tyrosinase-inhibitory activities, indicating that their bioactivities are not uniform but strongly influenced by species identity. (2) Several seed extracts showed complementary multifunctional properties, supporting their potential use as multi-target ingredients rather than single-purpose functional materials. (3) The utilization of preliminary-grade seeds provides a sustainable upcycling strategy that improves resource-use efficiency while minimizing additional harvesting pressure on natural forest ecosystems. (4) Although the bioactivities were evaluated using crude extracts, the results provide foundational evidence supporting the potential of forest tree seeds as raw materials for cosmetic, pharmaceutical, and health-related applications. Further studies focusing on the identification of key bioactive compounds and elucidation of their molecular mechanisms are required to enable the rational development of high-value products derived from forest tree seed resources.

